# Colorimetric and fluorescent TRAP assays for visualising and quantifying fish osteoclast activity

**DOI:** 10.1101/2021.12.10.472045

**Authors:** Lalith Prabha Ethiraj, Fong En Lei Samuel, Ranran Liu, Christoph Winkler, Tom James Carney

## Abstract

Histochemical detection of tartrate-resistant acid phosphatase (TRAP) activity is a fundamental technique for visualizing osteoclastic bone resorption and assessing osteoclast activity status in tissues. This approach has mostly employed colorimetric detection, which has limited quantification of activity *in situ* and co-labelling with other skeletal markers. Here we report simple colorimetric and fluorescent TRAP assays in zebrafish and medaka, two important model organisms for investigating the pathogenesis of bone disorders. We show fluorescent TRAP staining, utilising the ELF97 substrate, is a rapid, robust and stable system to visualise and quantify osteoclast activity in zebrafish, and is compatible with other fluorescence stains, transgenic lines and antibody approaches. Using this approach, we show that TRAP activity is predominantly found around the base of the zebrafish pharyngeal teeth, where osteoclast activity state appears to be heterogeneous.

## Introduction

Bone remodeling is a normal physiological process for maintenance of bone mineral density, which is accomplished by harmonized function of osteoclasts and osteoblasts. Osteoclasts are responsible for removal of bone, whereas the osteoblasts carry out new bone formation (1). Disruption of the balance between osteoclastic bone resorption and osteoblastic bone formation can be seen in various bone diseases such as Osteogenesis imperfecta, osteoporosis, metastatic cancer and Paget’s disease (2). Accordingly, observation of osteoclast number and activity *in situ* is a critical tool in assessing the pathogenesis of skeletal disease. Osteoclasts are generated from the myeloid lineage under the influence of Macrophage colony-stimulating factor (M-CSF) and its receptor, Csf1r (3, 4), whilst osteoclasts can be activated by binding of receptor for activation of nuclear factor kappa B (NF-kB) ligand (RANKL) to its receptor RANK (5, 6).

Once activated, osteoclasts express Tartrate resistant acid phosphatase (TRAP or TRAcP) which is involved in hydrolysis of bone matrix (7), and its activity can is used as a proxy of osteoclastic bone resorption activity (8). The phosphatase activity of TRAP forms the basis of its histological detection. Presence of the enzyme within a sample removes phosphate from an inert napthol AS-BI phosphate substrate, which then is able conjugate with a daizonium salt to form a red precipitate. This colorimetric reaction can thus be visualized by light microscopy. TRAP is resistant to tartrate inhibition, distinguishing it from alkaline phosphatases of other cells in vivo (8).

A fluorescence-based staining method has also been developed for detection of TRAP activity (9). This employs the ELF97 substrate which yields a brightly fluorescent precipitate upon removal of a phosphate group by a phosphatase. The dephosphorylated product is maximally excited at 360nm while emission occurs maximally at 530nm. Such a long Stokes shift reduces interference caused by autofluorescence or other stains and greatly expands ability to combine with other fluorescent stains (10, 11).

Fish species are increasingly being used as models of bone disease and to study the biology of the skeletal system, and they show a strong conservation of genetic control of bone homeostasis (12). Here, we describe protocols and applicability of colorimetric and ELF97 based TRAP assays in zebrafish and medaka, and show the versatility fluorescent based TRAP staining approach provides to visualizing osteoclast activity in fish.

## Materials And Methods

### Fish husbandry

Adult zebrafish were maintained in a recirculation system at the NTU Zebrafish Facility at 28°C with a 14-and 10-hour light and dark cycle. All investigations were carried out in accordance with approved IACUC protocols (A18002) at Nanyang Technological University. The AB wild type strain was used for most experiments, whilst the *csf1ra*^*j4e1*^ (*panther*) mutant strain was used to genetically deplete osteoclasts. Labelling of osteoclasts and osteoblasts employed the *ctsk:dsred; sp7:egfp* transgenic zebrafish (13). Transgenic *HS:rankl* medaka used were reared in the fish facility at the Department of Biological Sciences (National University of Singapore) under the IACUC protocols (BR19-0120, R18-0562). Medaka larvae were heat shocked at 9dpf in a 39°C water bath for 2 hours. All embryos were obtained from natural crosses and were raised in E3 medium at 28°C until 12dpf. Adult and larval fish were anesthetised using 0.02% Tricaine (Sigma A5040) buffered to pH7.0 with Tris. Fish were euthanised using immersion in 0.4% Tricaine (pH 7.0).

### Bisphosphonate treatment of Zebrafish

Adult fish were treated with 100μg/ml Alendronate solution obtained by dissolving Alendronate (Sigma A4978) in fish tank water. The 12dpf zebrafish larvae were treated using 40μg/ml Alendronate solution dissolved in E3 medium. For treatment of adults, fish were immersed in the drug solution for 24 hours once a week for 5 weeks. Following each exposure pulse, treated fish were placed in a tank of fresh tank water for 2-day recovery, and then returned to the flow racks until the next exposure cycle. Larvae were treated by single immersion in the drug solution for 14 hours immediately before experiments.

### Zebrafish Jaw and Pharyngeal teeth dissection

To extract the teeth bearing 5^th^ ceratobranchial arch, adults were euthanised and laid laterally on a petri dish under a dissecting microscope. Drummond number 5 Watchmaker’s forceps were inserted from a posterior angle under the operculum, and the bony structure just below the gills were extracted out together with the surrounding soft tissue. Tissues were lightly cleared away from the bone manually using Watchmaker’s forceps and Vannas microsurgical scissors. Specimens were then processed by staining and then placed on a glass slide for imaging under a Zeiss dissecting fluorescent or confocal microscope.

### Fish tail fin ray fractures

After anaesthetisation adult zebrafish were positioned laterally under a dissecting stereomicroscope with tail spread out. A paper tissue was used to remove excess water. A fin ray was fractured by crush injury at the centre of a ray segment using forceps (Drummond number 5) according to Tomecka, Ethiraj (14). Up to 4 crushes were made per fin on non-adjacent fin rays and care was taken not to impact the neighbouring tissues. After injury, fish were transferred to a quiet tank for brief recovery. After the stipulated crush healing duration, the whole tail fin was dissected using a scalpel under anaesthetic, isolated tail fin subjected to staining and imaging.

### Tissue fixation

Prior to staining, all tissues and larvae were fixed in 4% paraformaldehyde (PFA) (Sigma: P6148) in phosphate buffered saline (PBS) on a rotator for 40 minutes at room temperature. The 4% PFA was removed from the samples with 3 washes of PBSTriton (0.1% Triton-X-100 (Promega: H5141) in 1x Phosphate Buffered Saline (PBS)) for 5 mins each at room temperature, and then processed for TRAP staining.

### Colorimetric TRAP staining

The colorimetric TRAP staining protocol was adapted from Takahashi, Udagawa (15). Briefly, 5mg of Naphthol AS-MX phosphate (Merck: N4875) was dissolved in 0.5 ml of N, N’-dimethylformamide (Merck: D4451) to make a 10mg/ml Naphthol AS-MX phosphate “Solution A” stock. A “Solution B” stock solution of 50mM Sodium tartrate and 1.6mM Fast Red Violet LB was made by dissolving 0.575g of sodium L-tartrate dibasic dihydrate (Merck: 228729) in 50 ml of 0.1M sodium acetate buffer (adjusted to pH 5.0 with 100% Acetic Acid), and then adding 30mg Fast Red Violet LB salt (Merck: F3381).

Final TRAP staining solution was obtained by mixing 1 part Solution A to 100 parts Solution B together (e.g., 0.5ml Solution A was added to 50ml Solution B). This final solution can be stored for 1 month in the refrigerator.

Following fixation and washing, the final PBSTriton wash was replaced with 1ml of TRAP staining solution and samples incubated at room temperature for 3 hours in the dark. Samples were then washed with PBST (0.1% Tween in PBS) 3 times at room temperature and then re-fixed with 4% paraformaldehyde for 30min.

### Sample Bleaching

If samples needed to be bleached to remove pigment following colorimetric TRAP staining, they were washed 3 times in PBST for 5 mins each following re-fixing, and a bleach solution (0.5% KOH and 3% H2O2 final concentrations) added. Samples were bleached at room temperature for 20mins with tube lids left open. Bleach solution was removed with thorough washing in PBST. Stained samples were washed into 90% glycerol for imaging and long-term storage in the dark at 4°C.

### Fluorescence TRAP staining protocol

The fluorescent TRAP staining approach was adapted from Filgueira (9). Briefly, Fluorescent ELF97-TRAP Staining Solution of 50mM Sodium tartrate and 0.2mM ELF97 was made by dissolving 0.575g of sodium L-tartrate dibasic dihydrate in 50 ml of 0.1M sodium acetate buffer (pH 5.0), and then adding 20μl 5mM ELF97 stock solution (Thermo Fisher Scientific E6589). Following fixation and PBSTriton washing, the final PBSTriton wash was removed, and the samples then incubated in Fluorescent TRAP staining solution for 2h at room temperature. Specimens were washed 3 times with PBST for 5min each, followed by re-fixation with 4% PFA for 30min at room temperature. Samples were again washed 3 times in PBST before transferring to 90%Gycerol for imaging and storage, or for additional fluorescent staining approaches.

### Combined ELF97 fluorescence staining protocol

Generally, for all combinatorial ELF97 and fluorescent labelling, ELF97 staining was performed first, and the second staining approach initiated after the re-fixation step. For DAPI counter staining, samples were washed out of PFA with 3 5min washes of PBST, followed by 30min incubation in 5μg/ml DAPI (4′,6-diamidino-2-phenylindole) in PBST for 30 minutes. Samples were rewashed in PBST before transferring to Glycerol.

For calcified tissue staining, the above was repeated but DAPI was replaced with 0.01% Calcein (Sigma: C0875) in PBST or 0.01% Alizarin Red (Sigma: A5533).

For immunofluorescent staining following ELF97-TRAP, samples were washed 3 times in PBST after re-fixing or glycerol storage and then subjected to −20°C Acetone for 20min to permeabilise. Methanol must not be used. Following 3 washes in PBSTriton samples were incubated in Block solution 2-3hrs room temperature (4% BSA, 1%DMSO, 0.5% Triton-X-100 in PBS) and immunostaining performed as per (16, 17). Primary antibodies used were chicken anti-GFP (Abcam: Ab16901) at 1:500 and rabbit anti-Dsred (BD Biosciences: #632496) at 1:200. Secondary antibodies used were Alexa488 Goat anti-chicken (Invitrogen: A11039) at 1:1000 and Alexa568-goat anti-rabbit (Invitrogen: A11011) at 1:200. Lastly, following secondary antibody staining, specimens were washed 3 times in PBSTriton for 30 minutes each and imaged and stored under 90% glycerol at 4°C without exposure to light.

### Cryosectioning

For fluorescent TRAP staining, cryosectioning was performed after the completion of all staining protocols. Samples stained with ELF97, with or without other fluorescent stains, were washed from PBST into OCT (Tissue-Tek Optimal Cutting Temperature freezing compound; Sakura Finetek) and orientated in cryomolds (Tissue Tek Cryomold). These were flash frozen in liquid nitrogen or through storage in a −80°C freezer. 20μm cryosections were made using Leica Cryostat (CM 3050). Sectioned fluorescent samples on slides were dried for at least one hour at RT or on a 42°C heat block. They were then washed with PBS, re-dried overnight at room temperature, and then covered in Vectashield mounting medium under a coverslip and imaged. For colorimetric TRAP staining, cryosectioning was performed before the TRAP staining protocol. After PFA fixation, samples were washed in PBS at 4°C overnight, and then substituted in 5, 10, 15 & 30% sucrose/PBS solution, for 30 minutes each. Finally samples were exchanged into 30% sucrose:OCT at 1:1 for 30min and in 100% OCT overnight. Samples were mounted in cryomolds and frozen. 20μm cryosections were made on a Leica Cryostat, and the slides dried overnight. Prior to beginning the colorimetric TRAP staining, slides were washed in 100% Methanol for 10mins and then PBS, and then subjected to the colorimetric TRAP staining protocol.

### Imaging

Samples were visualised using widefield Colibri on an AxioImager M2 (Zeiss), or confocal fluorescent microscopy (Zeiss LSM800) and processed using Zen software (Zeiss) or Fiji (ImageJ, ver. 1.52p). For the ELF 97 crush injury time series assay, corrected total crush fluorescence was used to measure fluorescent signal emitted (14, 18). All measurements were normalised against fluorescence detected at an unfractured site. For ELF97 detection, samples were illuminated with 405nm wavelength laser, and the emission detected using a filter collecting between 515nm – 617nm. For the Colibri system, samples were illuminated with the 385nm LED, and the mCherry Filter Cube used as the emission filter.

### Statistics

Statistical analysis was performed using Prism GraphPad software. ANOVA with either Bonferroni or Tukey post-tests were used.

## Results

### A colorimetric TRAP staining approach to detect osteoclastic activity in zebrafish and medaka

We evaluated our colorimetric TRAP staining method using a zebrafish tail fin crush injury model (14). We saw that TRAP staining is exclusive to fracture sites at 4 days post crush (4dpc) and not in adjacent unfractured rays (Fig 1A). We demonstrated that this staining of fractured fin rays was indeed reporting osteoclastic TRAP activity as it was largely abolished in the osteoclast deficient mutant *csf1ra*^*j4e1*^ mutant (Fig 1B) (19, 20). Our colorimetric method also detected TRAP activity at the 5th ceratobrachial arch in 12dpf larvae, where specific osteoclast TRAP activity around the pharyngeal teeth is well known to occur (21, 22). No TRAP positive cells around the pharyngeal teeth regions were observed in the *csf1ra*^*j4e1*^ mutant as expected (13), nor was any staining noted in other pharyngeal arches of the WT, confirming the activity to be associated only with osteoclastic activity in teeth bearing arches of 12dpf larvae (Fig 1C-D).

**Figure 1:**
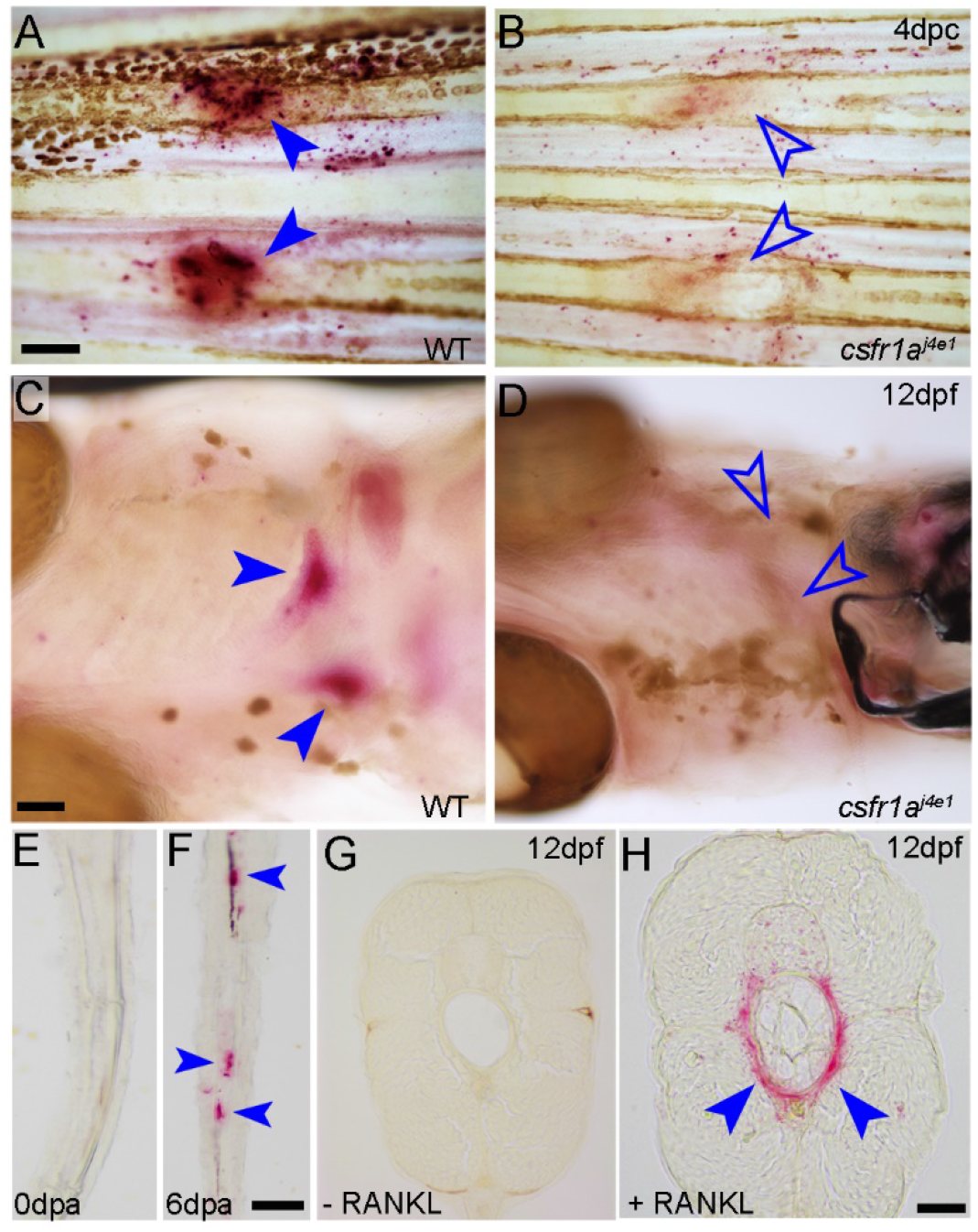
Identification of activated osteoclasts using conventional TRAP stain. **A-H:** Brightfield images of zebrafish (A-D) and medaka (E-H) adult fins (A-B, E, F), or 12dpf larvae (C, D, G, H) processed by colorimetric TRAP staining. Individuals are either wildtype (WT; A, C, E, F), homozygous *csf1ra*^*j4e1*^ mutants (B, D) or transgenic for *HS:rankl* (G, H). Images are of wholemount fins 4 days following crush (4dpc; A, B), wholemount ventral views (C, D) or longitudinal (E, F) or transverse cryosections (G, H). Fins in (E, F) have been amputated and stained immediately (0dpa; E) or after 6 days (ddpa; F). Larva in (H) has been subjected to heatshock to express RANKL, while larva in (G) was not heatshocked. Red precipitate (blue arrowheads) indicates TRAP locations at fractures (A), pharyngeal teeth of the 5^th^ ceratobranchial arch (C), in regenerating fin rays at d days post amputation (6dpa; F) or around notochord in larvae with excess RANKL expressed by heatshock (H). TRAP staining is absent when there is lack of RANKL overexpression (G), at the start of fin regeneration (0dpa; E) or genetic ablation of osteoclasts (B, D; open blue arrowheads), indicating specificity. Scale bars: (A) = 200μm, (C) = 100μm, (F, H) = 50μm.

Finally, we wanted to confirm the applicability of our staining protocol to other fish species. We used medaka which had osteoclast activity primed by two methods. Firstly, we amputated the adult tail fin and performed colorimetric TRAP staining on cryosections either immediately (0dpa) or after 6 days of regeneration (6dpa). TRAP staining was found in the blastema region, near segments only after 6 days, suggesting that osteoclasts were active in the regenerating fins only after regeneration had progressed (Fig. 1 F). Secondly, we used a transgenic approach to activate osteoclasts at 12dpf through a heat-shock transgenic approach to conditionally over-expresses RANKL (23). 9dpf *HS:rankl* medaka larvae were heatshocked for 2h at 39°C and TRAP staining performed on cryosections from the trunk 3 day later. *HS*:*rankl* larvae showed high osteoclast activity around the notochord (Fig 1G), but this was absent without heatshock (Fig 1F), in line with the previously shown increase in ectopic osteoclast numbers at this site following RANKL induction (23). Thus, the colorimetric TRAP staining approach offers a simple cheap method for defining osteoclast activity in fish species in both wholemount and following cryosections.

### Application of a fluorescent TRAP staining approach to zebrafish

Zebrafish are highly amenable to fluorescent microscopy approaches. We tested to see if a described fluorescent TRAP staining approach could be adapted to zebrafish. We tested the ELF97 substrate initially in the tail fin crush injury model. As with the colorimetric assay, we observed a strong fluorescent signal 3dpc in fractured rays only and not in adjacent intact rays (Fig 2A). We again showed that this signal was osteoclast dependant as it was absent in *csf1ra*^*j4e1*^ mutants and in fish treated for 14h prior to crushing with 100μg/ml of the bisphosphonate (BP), alendronate, which acts as an osteoclast poison (Fig 2B, C). We followed a time course of healing under the three conditions and saw that TRAP staining persisted in the WT crush at 7, and 14dpc but had largely resolved by 21dpc (Fig 2D, G, J), whilst it was mostly absent at all stages in the two osteoclast deficient scenarios (Fig 2E, F, H, I, K, L). Being a fluorescent stain, ELF97 facilitated quantification of TRAP activity *in situ* (Fig 2M). Mean fluorescence detected in *csf1ra*^*j4e1*^ mutants and BP fractures were noted both to be significantly reduced at 4, 7 and 14dpc compared to WT (Fig 2A-I, M). At 21dpc, fluorescence in WT was significantly reduced compared to the previous timepoint, while mean fluorescence in *csf1ra*^*j4e1*^ mutant fractures was significantly more than following BP treatment (Fig 2A, D, G, J, M). Statistical trends observed are in agreement with previous observations of osteoclast activity in fractures (14). However, ELF97 fluorescent imaging made quantification far less subjective and markedly improved analysis throughput.

**Figure 2:**
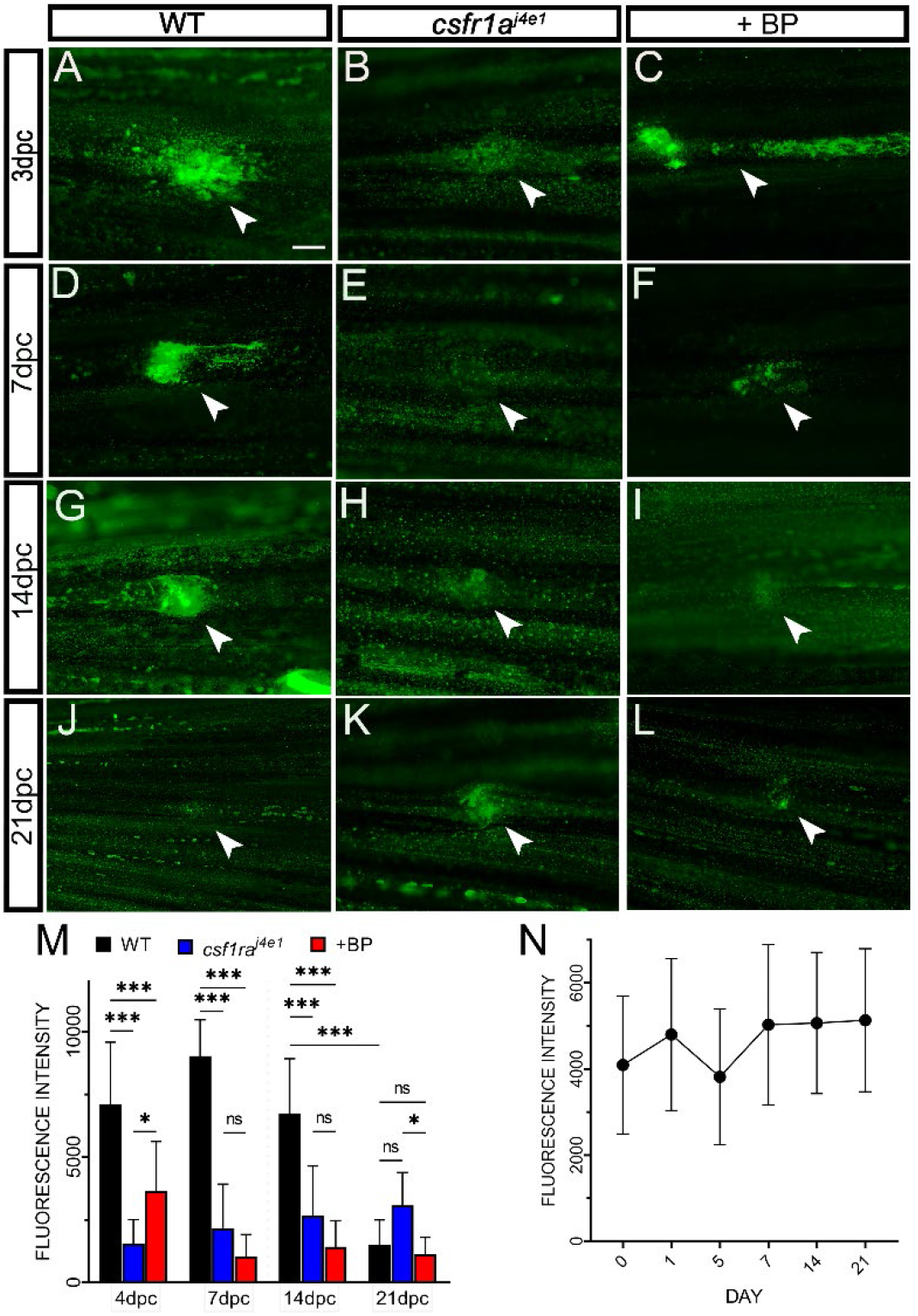
A fluorescent TRAP method labels osteoclast activity in zebrafish bone fractures. **A-L:** Wholemount widefield fluorescent images of adult fin ray fractures processed for ELF97 fluorescence. Fins were fixed at 3 days post crush (3dpc; A-C), 7dpc (D-F), 14dpc (G-I) and 21dpc (J-L), and were either WT (A, D, G, J), homozygous *csf1ra*^*j4e1*^ mutants (B, E, H, K) or treated with 100μg/ml Bisphosphonate. (C, F, I, L). **M**: Mean fluorescence of crush sites at given timepoints and under the three conditions. ***= p<0.001, *=p<0.05, ANOVA with Bonferroni multiple comparisons test, n=12. **N**: Mean ELF97 fluorescence at the crush sites of 1dpc WT zebrafish following storage for 0, 1, 5, 7, 14, 21 days. There was no statistically significant variation over the course of 21 days of storage compared against initial staining. ANOVA, Tukey’s multiple comparisons test, n=12. Scale bar (A) = 100μm.

To determine if ELF97 stained samples can be imaged at later dates, we assessed the stability of TRAP fluorescence reaction product after storage. WT tail fins which had been fractured were fixed after 1 day and processed using the ELF97 TRAP staining protocol. They were then stored in the dark at 4°C under glycerol and imaged repeatedly using confocal microscopy at timepoints 0, 1, 5, 7, 14 and 21 days after staining. The mean fluorescence of TRAP staining at the fractures showed no statistically significant change in intensity over the 21 days of storage, indicating stability of ELF97 fluorescence (Fig 2N).

### Fluorescent TRAP staining is compatible with fluorescent protein imaging and fluorescent nucleii and bone staining

Fluorescent microscopy provides the ability to image multiple fluorescent labels simultaneously, facilitating co-localisation studies. The large Stokes shift of ELF97 obviates interference with standard fluorophore spectra. Thus, we assessed if we could combine ELF97 TRAP staining with well used transgenic markers of the zebrafish skeletal system. In particular, the *cathepsin K (ctsk)* gene is expressed in osteoclasts, whilst the *sp7 (= osterix*) transcription factor is expressed in osteoblasts. The promoters of these genes have been used to label these cells with fluorescent proteins in transgenic fish. We performed tail fin ray fractures on *ctsk:dsred; sp7:egfp* double transgenic adults. At 7dpc, tails were briefly fixed and processed for ELF97 TRAP staining and then counterstained with DAPI. Confocal microscopy of the fracture site of the fin rays revealed that the ELF97 fluorescence co-localised within the field of osteoclast-specific DsRed fluorescence, which appeared at the fracture site (Fig. 3A, B, D-F). Osteoblast-specific green fluorescent protein (GFP) was noted most prominently in the peripheral bone but also showed localised staining within the fracture (Fig. 3C).

**Figure 3:**
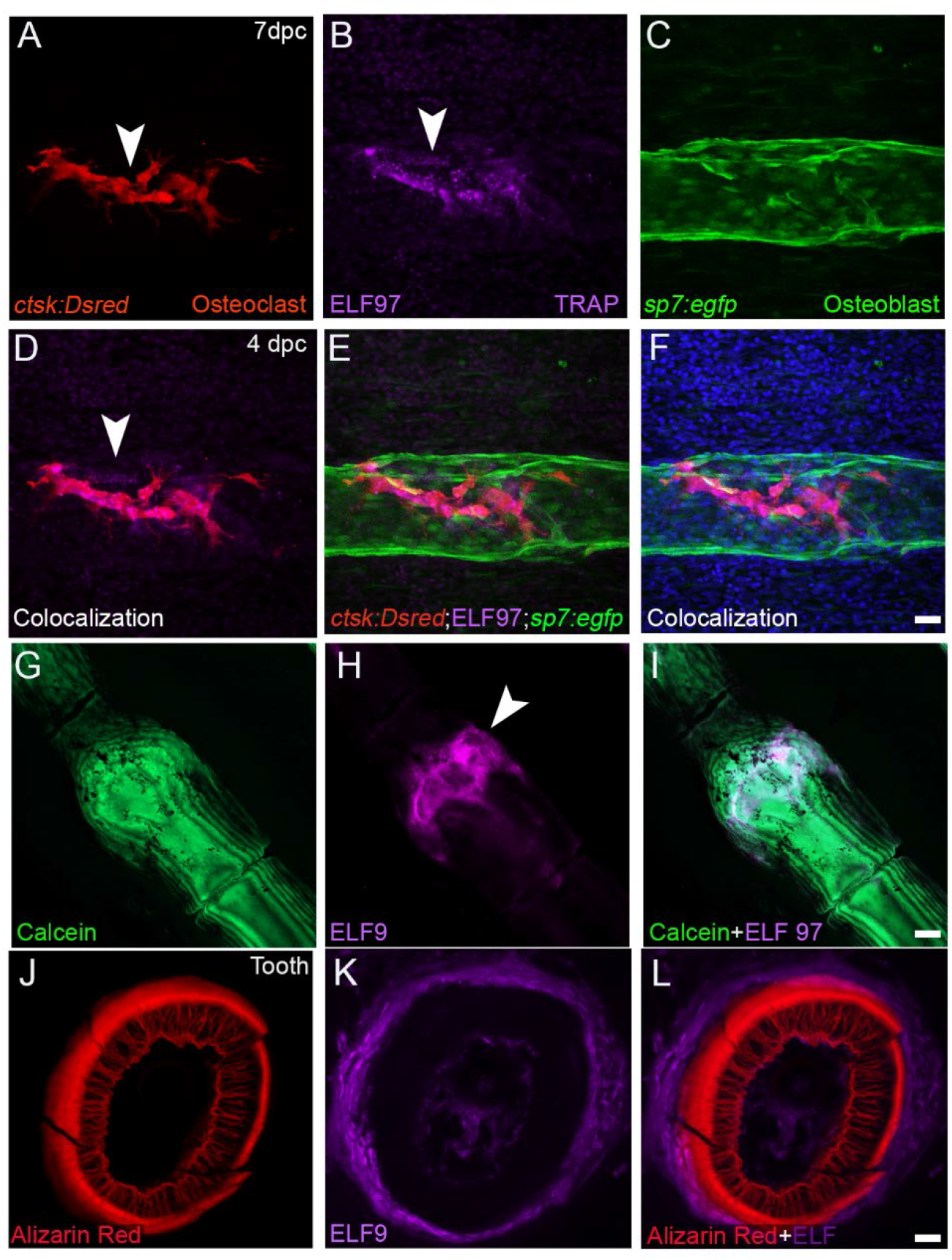
Colocalization of TRAP-ELF97 reaction product with fluorescent proteins and dyes. **A-F:** Wholemount confocal images of a fin ray fracture in *ctsk:Dsred; sp7:egfp* double transgenic adult, processed at 7dpc for ELF97 TRAP fluorescence and counterstained with DAPI. Individual fluorescent channels for *ctsk:Dsred* (A), ELF97 (B), *sp7:egfp* (C), DsRed and ELF97 channel overlay (D), DsRed, EGFP and ELF97 channel overlay (E), all channels including DAPI (F). TRAP-ELF97 fluorescence is seen associated with DsRed labelled osteoclasts within the callus. **G-I:** Confocal images of wholemount adult zebrafish fin ray subjected to crush fracture and processed at 4dpc for Calcein (G) and ELF97 (H) staining. TRAP activity is seen within the bone matrix of the callus (I). **J-L:** Confocal image of cryosection through an adult zebrafish pharyngeal tooth which had been stained with Alizarin Red (J) and ELF97 (K). Overlay of both channels indicates TRAP staining surrounding the bone mineral (L). Scale Bars: (F, I, L) = 20μm

To assess if our fluorescent TRAP staining approach was compatible with fluorescent bone stains, we used ELF97 in combination with Calcein in whole-mount staining of adult fin ray fractures (Fig 3G-I) and with Alizarin red in cryosections of the adult pharyngeal teeth where there is continual osteoclast activity (Fig 3J-L). In the former, ELF97 staining was seen again within parts of the fracture callus, whilst in the cryosections, calcified enamel lamellae of the teeth were circumscribed by osteoclast active regions.

### ELF 97 allows combination with immunofluorescent staining

In many samples, residual fluorescence from the *ctsk:dsred; sp7:egfp* double transgenes was not visible following TRAP staining. In these cases, it was necessary to detect DsRed and EGFP by immunostaining. To assess if the ELF97 protocol was indeed compatible with immunostaining. We isolated the 5th ceratobrachial arch of *ctsk:dsred; sp7:egfp* double transgenic adults by dissection and fixed overnight in 4% PFA. Following this, there was no observable DsRed or EGFP fluorescence remaining. We performed ELF97 TRAP staining on the samples and then immunofluorescently labelled them using antibodies against DsRed and EGFP. We imaged them in wholemount and additionally cryosectioned them to ascertain if the labels remained. We saw that the TRAP staining was not disrupted by the immunofluorescent protocol, and that ELF97 was compatible with antibody detection of DsRed and EGFP, both in wholemount (Fig 4A-F) and in cryosections (Fig 4G-M). EGFP signal could be seen in the pulp, dentine and bone of attachment, previously shown as expression domains of *sp7* (Fig 4A, E, F, G, J, L, M) (24). The expression domain of EGFP was quite distinct from that corresponding to DsRed, which was predominantly found in the bone regions below each tooth, and around the base indicating distinct localisation of osteoclasts and osteoblasts (Fig. 4B, D, E, I, K-M). ELF97 signal was found in a domain largely overlapping the DsRed domain, although the overlap was not identical, and there were clear locations where osteoclasts were not positive for the TRAP stain and others where the TRAP stain was broader than the DsRed staining (Fig. 4B, D, I, K). Thus, it appears there is heterogeneity of osteoclast state around the base of each tooth in zebrafish, and on the arch bone itself.

**Figure 4:**
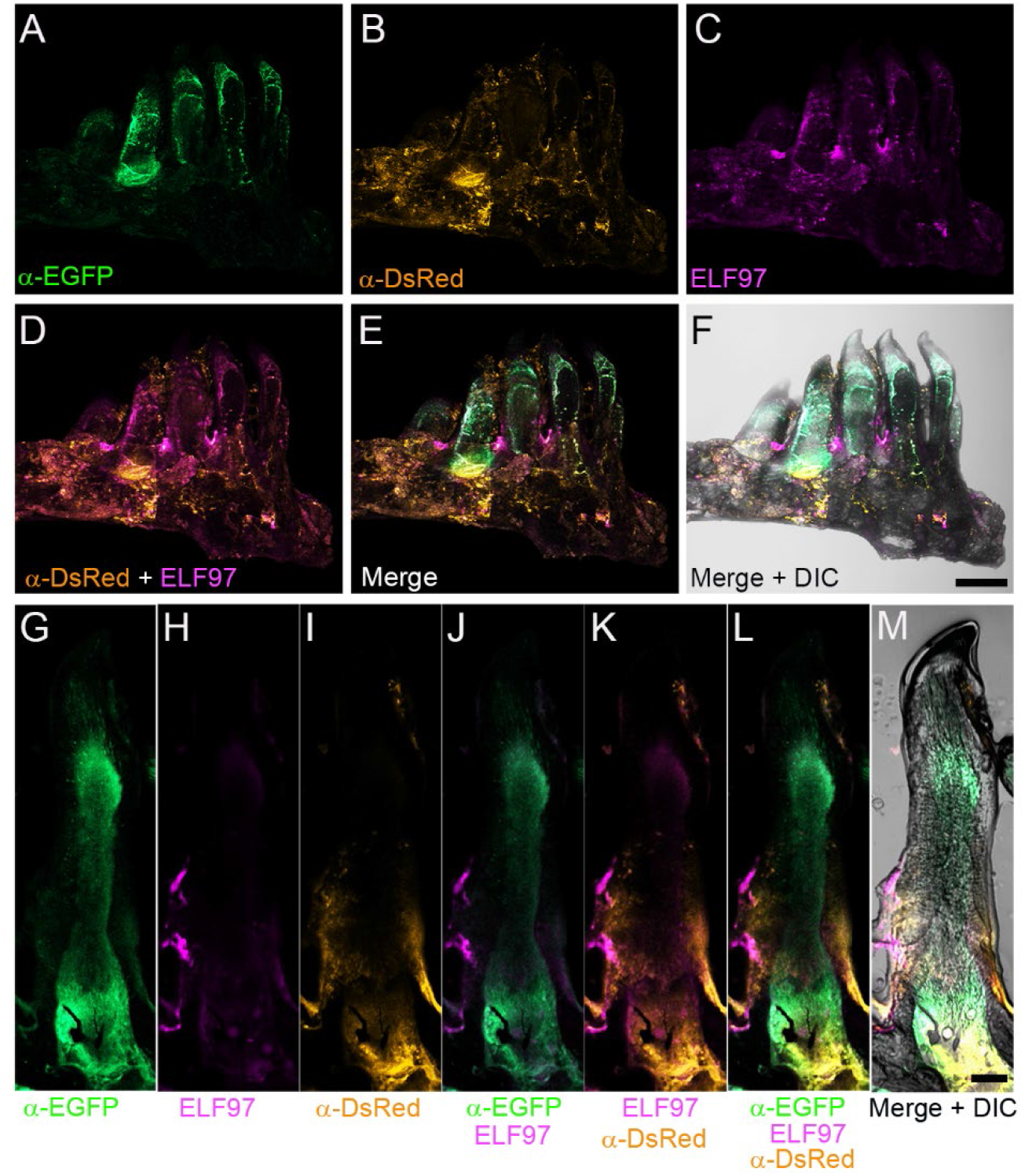
Fluorescent TRAP staining can be combined with immunofluorescence. **A-M:** Confocal images of dissected 5^th^ ceratobranchial arch with pharyngeal teeth of a *ctsk:Dsred; sp7:egfp* double transgenic adult which had been processed for ELF97 TRAP staining (C, H) and immunostained with antibodies against EGFP (A, G) and DsRed (B, I). Both wholemount (A-F) and cryosectioned (G-M) samples are presented. Co-localisation demonstrated ELF97 signal associated with DSRed positive osteoclasts found at the base of the teeth (D, E, K, L), whilst eGFP positive osteoblasts are found within the teeth (F, J, L). Fluorescent channels are also shown with DIC image (F, M). Scale bars: (F) = 200μm, (M) = 50μm.

## Discussion

We have thus shown that ELF 97 offers a simple, reproducible, stable, versatile, and quantifiable substitute to conventional colorimetric TRAP stains for analysis of fish osteoclast activity. It can be combined with other fluorescent dyes, mineral stains, fluorescent proteins and immunofluorescence, and is compatible with both wholemount and cryosectioning. Since zebrafish is a polyphyodont (25), there is continuous shedding of teeth throughout their life cycle. This provides an excellent model to study spontaneous continuous osteoclastic activity, which can be demonstrated using the ELF97 fluorescent stain. Combining this stain with other fluorescence staining approaches, we have identified that TRAP activity is not entirely concordant with presence of osteoclasts in the tooth region or in fractures, but appears to only partially overlap. This highlights the utility of assessing osteoclast activity in conjunction with their location and morphology. With fish systems providing increasingly sophisticated models of skeletal diseases, such methodologies will complement the growing array of genetic and labelling resources available.

## Acknowledments

We thank Dr. Matthew Harris, Department of Genetics, Harvard Medical School, Boston for gifting us the *ctsk:dsred; sp7:egfp* double transgenic line.

## Notes

### Competing Interest Statement

The authors have declared no competing interest.

